# Exposure to childhood adversity and adult alcohol, nicotine, and illicit drug use in Germany: a cross-sectional analysis of the German National Cohort (NAKO)

**DOI:** 10.64898/2026.10.12.758429

**Authors:** Justin Böhmer, Kanya Meyer-Degering, Ali Ghadami, Abhijit Sreepada, Patricia Bohmann, Stefanie Castell, Manon Feuchtinger, Cornelia Gottschick, Hans J. Grabe, André Karch, Claas Lahmann, Alexander Lang, Michael Leitzmann, Claudia Meinke-Franze, Dörthe Meyerdierks, Rafael Mikolajczyk, Katharina Nimptsch, Nadia Obi, Annette Peters, Tobias Pischon, Michael J. Stein, Thomas Keil, Lilian Krist, Heike Tost, Christine Heim

## Abstract

**Rationale:** Exposure to adversity in childhood and adolescence (i.e., “early-life stress”; ELS) is a well-established risk factor for substance use.

**Objectives:** The aim of this study was to investigate associations between ELS exposure and substance use across subtypes of ELS, different substance classes, and behavioral profiles of substance use.

**Methods:** In this cross-sectional study, we examined associations between ELS exposure and substance use in a population-based sample of 172,173 adults aged 19-74 years from the German National Cohort (NAKO). ELS exposure was assessed using the Childhood Trauma Screener. Substance use included nicotine, alcohol, and illicit drug use. Associations were examined using regression models evaluating overall ELS exposure, cumulative trauma load, and ELS subtypes. Latent class analysis (LCA) was conducted to identify substance use profiles.

**Results:** ELS exposure was associated with higher odds of smoking, greater smoking severity, heavier alcohol consumption among drinkers, and higher odds of lifetime illicit drug use. Overall, the strongest associations were observed for opioid and stimulant use. Dose–response analyses revealed higher risk with increasing numbers of ELS subtypes. Abuse-related subtypes, particularly exposure to physical and sexual abuse, consistently showed strongest independent associations across substances, whereas neglect exhibited weaker, more variable associations. LCA identified four substance use profiles, with ELS exposure predicting membership in more severe polysubstance classes.

**Conclusions:** ELS emerged as a robust risk factor for adult substance use, underscoring the need for trauma-informed approaches in clinical practice, public health, and policy to mitigate the long-term consequences of early adversity.

## INTRODUCTION

Childhood adversity or early-life stress (ELS), including sexual, physical, and emotional abuse, and/or physical and emotional neglect, among other stressors experienced during childhood or adolescence, is highly prevalent (Madigan et al., 2026), with around 60% of individuals having experienced at least one adverse childhood experience (Madigan et al., 2023). ELS is a pervasive risk factor for adverse mental and physical health outcomes across the lifespan (Heim et al., 2019) [3]. Extensive evidence confirms that exposure to ELS substantially increases risk for a wide range of psychiatric disorders and physical health conditions, constituting a major public health problem (Shonkoff et al., 2012; Teicher et al., 2022).

Prior research provides mechanistic insights into the link between ELS exposure and subsequent health problems. Compelling evidence suggests that ELS exposure induces profound and lasting alterations of multiple regulatory systems during sensitive developmental periods, resulting in enduring shifts in physiological and neural setpoints that shape stress responsivity and behavioral regulation across the lifespan (Heim et al., 2019). In particular, ELS exposure has been associated with long-term alterations in hypothalamic–pituitary–adrenal (HPA) axis function, as well as structural and functional changes in neural circuits supporting stress responsivity, reward processing, and executive control (Heim et al., 2000; Lupien et al., 2009; Teicher et al., 2016). Such alterations are hypothesized to bias individuals toward heightened stress sensitivity, altered reward valuation, and reduced regulatory control — core processes implicated in the development and maintenance of addiction-related behaviors (Koob & Volkow, 2016). In this regard, individuals exposed to early adversity are more likely to initiate substance use at younger ages (Dube et al., 2003), develop more severe patterns of use (Larance et al., 2018), and experience poorer treatment outcomes and higher relapse rates (Van Dam et al., 2014). These associations have been reported across multiple substance categories, including alcohol, tobacco, and illicit drugs, suggesting that ELS exposure may confer a broad vulnerability to substance-related behaviors (Enoch, 2011; Hughes et al., 2017; Leri et al., 2026).

Despite this substantial body of evidence, important gaps of knowledge remain: First, much of the existing literature on the association between ELS exposure and substance use in adulthood is based on relatively small or selected samples, limiting statistical power and generalizability. Second, ELS exposure is commonly operationalized using cumulative trauma scores. While informative, this approach may obscure the differential impact of specific trauma subtypes, such as abuse- or neglect-related forms, on later substance use. Subtypes characterized by threat are believed to have especially pronounced effects on neural systems supporting emotional and reward processing, relative to deprivation-related subtypes (Sheridan & McLaughlin, 2014), and might therefore be particularly influential for substance use behavior. Third, although individuals with a history of ELS exposure are at greater risk for polysubstance use (Clark et al., 2023), much less is known about the specific patterns of substance use behavior they may show. Individuals often engage in complex combinations of use across different substances, which may reflect distinct behavioral, psychological and neurobiological profiles. However, most prior studies have treated substance use variables independently, failing to capture the heterogeneity and co-occurrence of substance use behaviors.

The present study addresses these limitations by leveraging data from the largest available population-based sample of adults across Germany. Specifically, we aimed to examine associations between ELS and adult substance use by assessing the effects of overall ELS exposure, investigating dose–response patterns between the number of experienced ELS subtypes and substance use outcomes, and evaluating subtype-specific associations between distinct forms of ELS and the use of a broad range of legal and illicit substances.

Based on prior research, we hypothesized that individuals exposed to moderate-to-severe ELS would show higher likelihood and severity of substance use compared to those reporting no or mild ELS exposure, with risk increasing with the number of ELS subtypes experienced, and elevated risk for abuse-related subtypes compared with neglect-related forms. To address the limited understanding of polysubstance use behavior, we further aimed to identify latent substance use profiles and test whether ELS exposure predicts differential class membership.

## METHODS

### Study design and participants

This cross-sectional study is based on data from the German National Cohort (NAKO), a population-based prospective study examining disease etiology and risk factors for a range of common diseases. Between 2014 and 2019, over 205,000 individuals aged 19-75 years were examined across 18 study centers in Germany. The data analyzed here comprised information from NAKO’s baseline assessment including 204,648 participants. A detailed description of the study design and procedures was provided elsewhere (Peters et al., 2022). The study protocol was approved by the ethics committees of all participating study centers. Written informed consent for study participation and data use was obtained from all participants. All study procedures and analyses were conducted in accordance with ethical regulations.

### Procedures

Sociodemographic information, including age, sex, educational attainment, and parental education and employment, was collected using a standardized computer-assisted face-to-face interview. Educational attainment was assessed based on years of education according to the International Standard Classification of Education (ISCED-97). Parental education was assessed on a 5-point Likert scale. A composite parental education score (range: 2–10) was calculated by summing maternal and paternal education scores. Parental employment captured each parent’s employment status when the participant was 13 years old and was coded dichotomously (employed vs. not employed). A composite parental employment score (range: 0–2) reflected the number of employed parents. Race and ethnicity were not examined as they were not part of the prespecified analytical framework for this study.

Exposure to ELS was assessed using the Childhood Trauma Screener (CTS), a five-item self-report measure covering emotional and physical neglect, and emotional, physical, and sexual abuse. For each ELS subtype, severity was dichotomized into no/mild versus moderate/severe ELS based on established cut-offs (Glaesmer et al., 2013). Participants meeting criteria of moderate/severe ELS exposure for at least one subtype were classified as having experienced early-life stress (ELS+). Additionally, depressive and anxiety symptoms were assessed using the Patient Health Questionnaire-9 (PHQ-9) and Generalized Anxiety Disorder Scale–7 (GAD-7), respectively. For substance use assessment, smoking-related measures were obtained via self-report. To assess lifetime occurrence of tobacco smoking, we grouped current smokers and former smokers into a single category of ever smokers. In addition, age at smoking initiation, lifetime smoking duration, and lifetime average number of cigarettes smoked per day were assessed. Alcohol consumption was assessed via self-report using the short form of the Alcohol Use Disorder Identification Test (AUDIT-C), capturing alcohol use frequency, quantity, and frequency of binge-drinking. Moreover, average daily alcohol intake (g/day) was calculated based on self-reported frequency and quantity of consumption of different alcoholic beverages over the past 12 months. To evaluate lifetime occurrence of illicit drug use, subjects were asked to indicate whether they had ever used one of the following substances: cannabis, synthetic cannabinoids (e.g., Spice, Smoke, Space), amphetamine-type stimulants, methamphetamine (crystal meth), 3,4-methylenedioxymethamphetamine (MDMA), lysergic acid diethylamide (LSD), heroin, non-heroin opioids (e.g., codeine, methadone, opium, morphine), powder cocaine, and crack cocaine.

### Statistical Analysis

We examined associations between ELS exposure and substance use based on a three-step analytic approach. First, we tested whether ELS exposure per se was associated with substance use by modeling exposure to any moderate-to-severe ELS as a binary predictor (i.e., ELS+ vs. ELS-). Second, we examined dose–response relationships by testing whether higher trauma load, defined as the number of subtypes with moderate-to-severe ELS, was associated with higher substance use. Third, all five ELS subtypes were entered simultaneously into multivariable models, allowing estimation of subtype-specific effects. Logistic regression models were fit for categorical substance use outcomes, and linear regression models were used for continuous outcomes. All models were adjusted for age, sex, site, educational attainment, parental education, and parental employment status. Moreover, we tested interactions between ELS and sex or age (modeled in 10-year increments) and examined depressive and anxiety symptoms as potential mediating pathways. Given the large sample size, even minimal effects are likely to reach statistical significance. Therefore, our interpretation focuses primarily on effect sizes. Analyses were performed using R (v4.4.2).

To account for the heterogeneity of substance use behaviors, we conducted latent class analysis (LCA) based on binary indicators of lifetime use across all investigated substances. Models with one to eight classes were estimated using the poLCA package in R. Model selection was guided by the Bayesian Information Criterion (BIC), with lower values indicating better model fit. To ensure model stability, each candidate model was estimated using multiple random starting values, and convergence to a consistent maximum likelihood solution was verified. Associations between ELS indicators and latent classes were examined using multinomial logistic regressions, adjusted for the same set of covariates as outlined above.

## RESULTS

### Sample Characteristics

The final sample for the present analysis comprised 172,173 participants (84.13% of total sample) with valid ELS data, of whom 45,190 subjects (26.25%) reported at least one moderate-to-severe ELS exposure. Physical neglect was the most frequently reported subtype. Sample sizes varied slightly across analyses (Table 1) due to partially missing data (Table S1).

### Nicotine Use

Exposure to any form of moderate-to-severe ELS was associated with higher lifetime odds for smoking, longer lifetime smoking duration, higher lifetime daily cigarette consumption, and an earlier age at smoking initiation (Table 2). Dose–response relationships were observed, with higher number of ELS subtypes being associated with greater risk for all smoking-related outcomes (Table S2). Among maltreatment subtypes, those characterized by abuse showed stronger independent associations with nicotine use than neglect-related forms, with physical abuse conferring the greatest risk (Figure 1).

**Figure 1.**
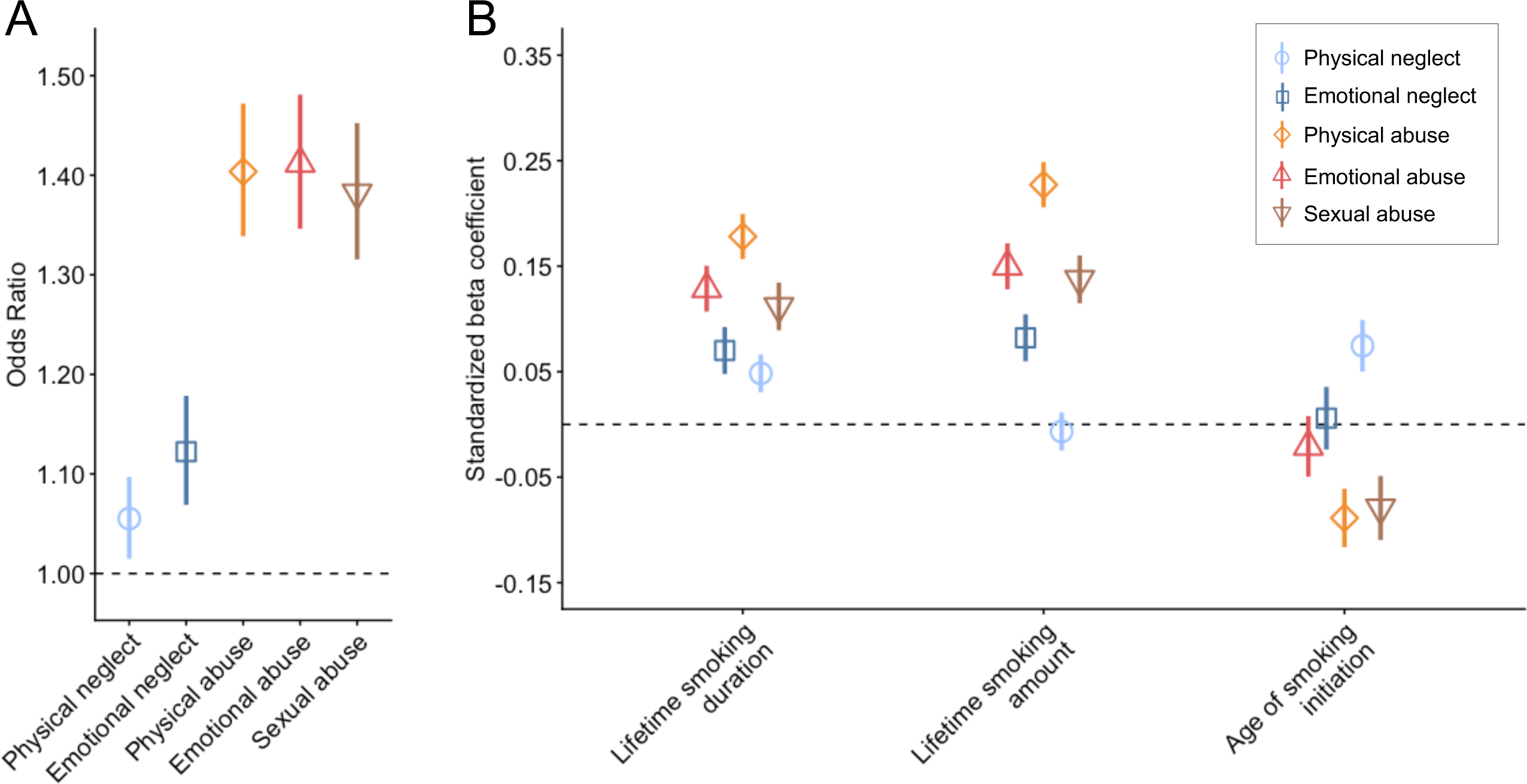
Association between early life stress (ELS) subtypes and smoking behavior. A) Odds ratios and 95% confidence intervals from logistic regression models examining the association between each ELS subtype and lifetime smoking occurrence. B) Standardized beta coefficients from linear regression models assessing the associations between ELS subtypes and continuous smoking outcomes, including lifetime smoking duration, lifetime smoking amount, and age of smoking initiation.

### Alcohol Use

Linear regression showed a small negative association between ELS exposure and AUDIT-C total score (Table 2). To account for potential non-linear associations across drinking levels, we applied multinomial logistic regression models categorizing abstinence (AUDIT-C = 0), moderate drinking (AUDIT-C 1-6 for females, 1-7 for males), and high-risk drinking (AUDIT-C ≥ 7/8 for females/males). ELS exposure was associated with higher odds of both abstinence (OR = 1.43, 95% CI [1.37, 1.50]) and high-risk drinking (OR = 1.22, 95% CI [1.15, 1.30]) relative to moderate drinking, indicating a polarization rather than a uniform decrease in alcohol use.

Further analyses among current drinkers showed that ELS exposure was associated with less frequent drinking but higher consumption when drinking, reflected in more drinks per occasion, more frequent binge-drinking episodes, and greater average daily intake (Table 2). Dose–response associations were observed for drinking frequency, drinks per occasion, and binge-drinking frequency (Table S2).

Across ELS subtypes, broadly consistent patterns emerged (Figure 2). All subtypes were associated with higher odds of abstinence relative to moderate drinking, with the strongest effects for emotional neglect and emotional abuse. Associations with high-risk drinking were weaker and more variable, showing modest strength for physical, emotional, and sexual abuse, but no association for neglect. For continuous drinking outcomes, all subtypes except sexual abuse were linked to lower drinking frequency. Physical abuse was the only subtype consistently associated with heavier consumption.

**Figure 2.**
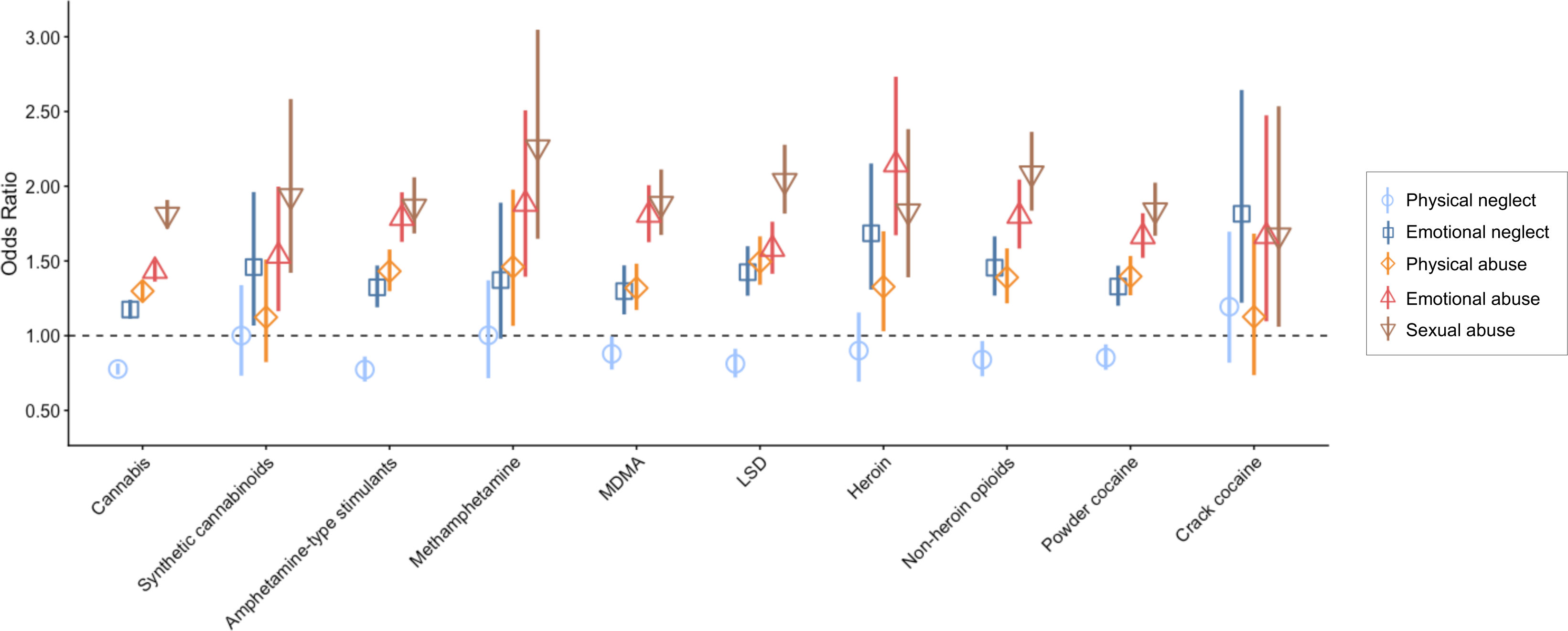
Association between early life stress (ELS) subtypes and alcohol drinking. A) Odds ratios and 95% confidence intervals from multinomial logistic regression models examining associations between each ELS subtype and alcohol use categories (abstinence and high-risk drinking) relative to moderate drinking. B) Standardized beta coefficients derived from linear regression models examining the associations between ELS subtypes and continuous drinking outcomes, including frequency of drinking, number of drinks per occasion, frequency of binge-drinking, and average amount of daily alcohol intake.

### Illicit Drug Use

ELS exposure was associated with higher odds of lifetime use across all illicit substances, with the strongest effects for heroin, methamphetamine, and non-heroin opioids (Table 2). ELS exposure was also linked to a greater number of substances ever used (β = 0.174, 95% CI [0.162, 0.186]). Experiencing multiple types of ELS further increased the odds of use across all substances (Table S2), suggesting dose-response relationships. The largest increment was observed for heroin and methamphetamine, with each additional ELS subtype increasing the risk of use by 56%. The number of ELS types experienced was likewise associated with a greater number of different substances used (β = 0.084, 95% CI [0.079, 0.090]).

Across substances, abuse-related ELS subtypes showed stronger associations with illicit drug use than neglect-related forms. Sexual abuse exhibited the strongest overall associations with lifetime use of most substances (Figure 3) and with the number of substances ever used (Table S3), with two exceptions: emotional abuse showed the strongest association with heroin use, whereas emotional neglect was most strongly associated with crack cocaine use.

**Figure 3.**
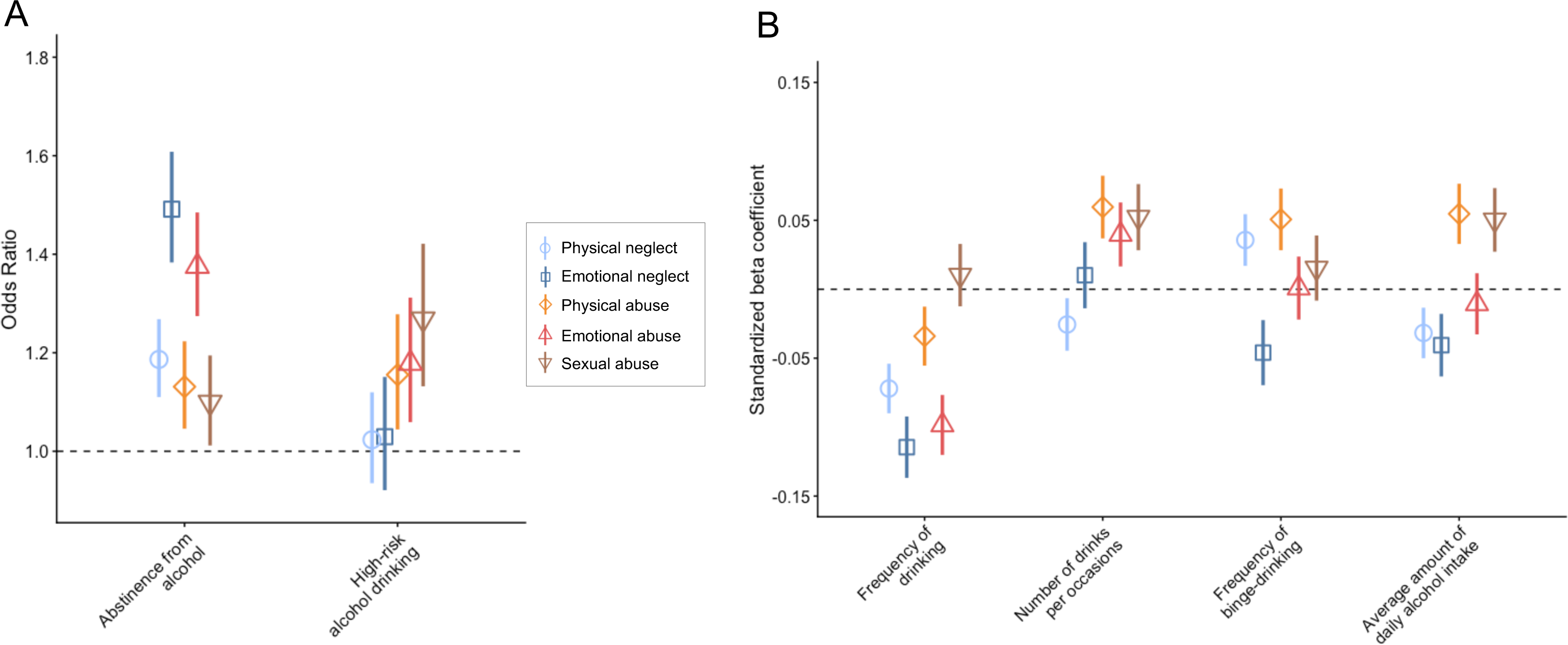
Odds ratios and 95% confidence intervals derived from logistic regression models examining the association between ELS subtypes and lifetime illicit drug use.

### Moderation and mediation analyses

Associations between ELS and substance use were generally stronger among younger individuals and, for several alcohol and illicit drug outcomes, among women compared to men (Supplementary Results; Table S4). Internalizing symptoms partially mediated these associations, accounting for 5– 63% of the total effects, with stronger statistical mediation via depressive than anxiety symptoms (Table S5).

### Latent Class Analysis of Substance Use Patterns

Latent class analysis yielded a four-class solution (Table S6) with stable convergence and well-differentiated substance use profiles (Figure 4A, Table S7). The largest class (75.22%; Figure 4A yellow line) showed minimal substance involvement with very low illicit use and modest nicotine/cannabis use (Class 1: *Minimal use*). A second class (20.30%; Figure 4A green line) showed elevated cannabis and nicotine use with little use of other substances (Class 2: *Cannabis–nicotine use*). A third class (4.09%; Figure 4A blue line) reflected frequent use of cannabis, nicotine, stimulants, and hallucinogens (Class 3: *Stimulant–hallucinogen use*). The smallest class (0.39%; Figure 4A red line) exhibited high probabilities across all substances, indicating a severe polysubstance pattern (Class 4: *High-risk polysubstance use*).

**Figure 4.**
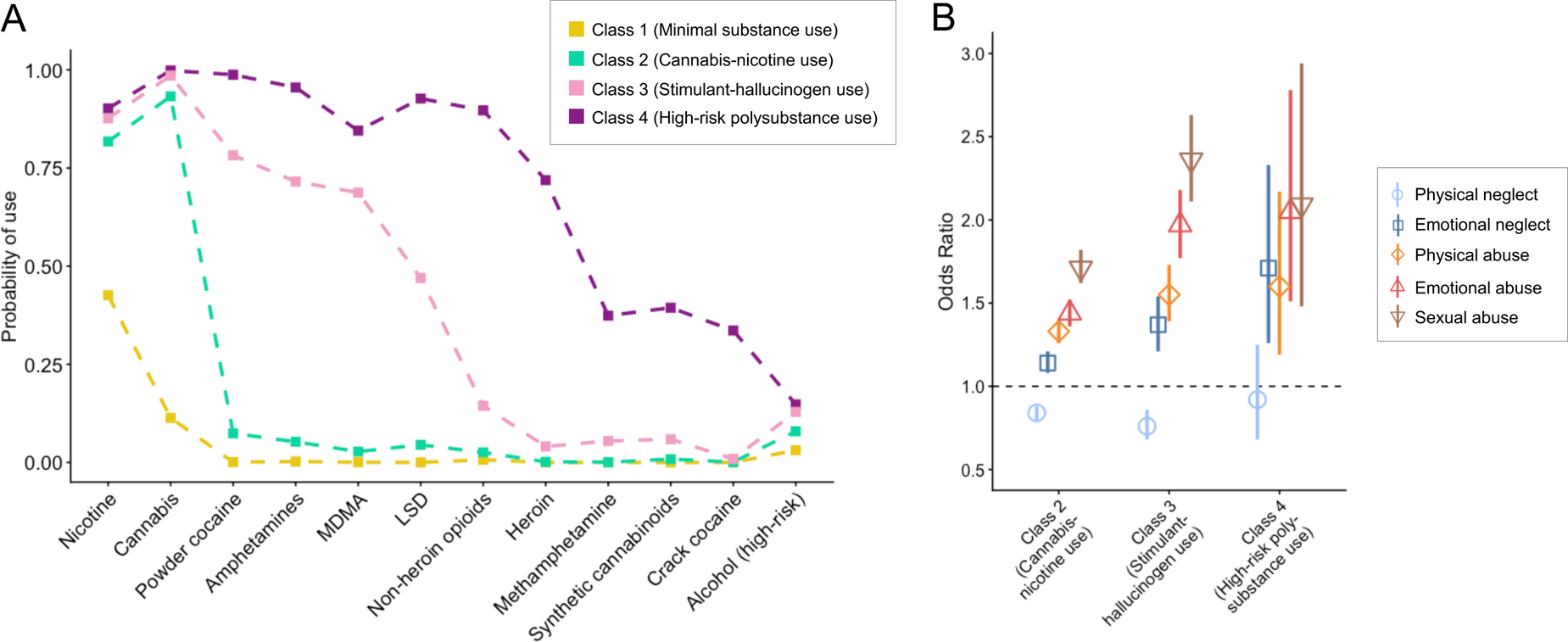
Latent class analysis (LCA) of substance use behaviors and association with early life stress (ELS) among 166,334 adults with complete data on all investigated substances. A) LCA identified four distinct substance use profiles, characterized by minimal use (Class 1; *n* = 125,110), predominantly cannabis and nicotine use (Class 2; *n* = 33,772), predominantly stimulant and hallucinogen use (Class 3; *n* = 6,804) and high-risk polysubstance use (Class 4; *n* = 648). Lines represent class-specific probabilities of use for each substance. (B) Odds ratios (ORs) and 95% confidence intervals from a multinomial logistic regression model estimating the association between ELS exposure and class membership. ORs indicate the relative odds of belonging to Classes 2-4 compared with the minimal-use reference class (Class 1).

Multinomial logistic regression analyses indicated a graded association between ELS exposure and substance use class membership. Compared with individuals from the *Minimal use* class, ELS exposure was associated with progressively higher odds of belonging to more severe substance use classes. Specifically, ELS exposure increased the odds of membership in the *Cannabis– nicotine* class by 46% (OR = 1.46, 95% CI [1.42, 1.51]), doubled the odds of membership in the *Stimulant–hallucinogen* class (OR = 2.04, 95% CI [1.92, 2.18]), and was associated with a more than twofold increase in the odds of belonging to the *High-risk polysubstance* class (OR = 2.32, 95% CI [1.91, 2.82]).

Experiencing a greater number of ELS types was associated with higher odds of belonging to any substance use class relative to the *Minimal use* class. Specifically, each additional ELS type was linked to a 26% increase in the odds of membership in the *Cannabis–nicotine* class (OR = 1.26, 95% CI [1.24, 1.28]), a 54% increase for the *Stimulant–hallucinogen* class (OR = 1.54, 95% CI [1.48, 1.59]), and a 64% increase for the *High-risk polysubstance* class (OR = 1.64, 95% CI [1.50, 1.79]).

Concerning ELS subtypes, sexual abuse demonstrated the strongest unique association with membership in any substance use class relative to the *Minimal use* class, conferring the highest relative risk among all ELS dimensions. In contrast, physical neglect did not show a positive association with substance use class membership and exhibited inconsistent patterns across classes (Figure 4B).

## DISCUSSION

The present study provides a comprehensive examination of the association between exposure to ELS and substance use in adulthood in a large, population-based cohort in Germany. Across substances, ELS exposure was consistently linked to elevated risk of substance use, more severe patterns of consumption, and greater likelihood of polysubstance involvement, with evidence for dose–response relationships. These findings extend prior work by demonstrating substance-specific patterns, with stronger associations for opioids and stimulants, as well as subtype-specific differences, with abuse-related forms of ELS showing the strongest effects relative to neglect-related forms. Together, these findings underscore the broad impact of ELS on adult substance use and point to distinct neurobiological and psychological pathways through which childhood adversity influences substance use vulnerability.

Consistent with previous research (Enoch, 2011; Hughes et al., 2017; Leri et al., 2026), exposure to ELS was associated with elevated odds of lifetime use across a broad range of legal and illicit substances. Moreover, our findings show that ELS exposure is linked not only to the likelihood of use but also to more severe patterns of consumption. Among individuals exposed to ELS, we observed dose-response relationships across all substance outcomes, supporting cumulative risk models of ELS (Putnam et al., 2013). The strongest associations for ELS exposure and cumulative adversity load were observed for heroin, non-heroin opioids, and methamphetamine. This pattern suggests that individuals with maltreatment histories are particularly vulnerable to substances with potent euphoric, anxiolytic, and affect-modulating properties. Mechanistically, exposure to ELS exerts enduring effects on neurobiological systems central to the reinforcing properties of opioids and stimulants. For instance, ELS exposure alters the development and function of the endogenous opioid system, heightening the rewarding effects of opioids while attenuating their aversive properties (Carlyle et al., 2021). Stimulants act predominantly on dopaminergic pathways, and evidence shows that individuals with exposure to ELS exhibit heightened dopaminergic responses following stimulant exposure (Oswald et al., 2014), suggesting increased sensitivity to their euphoric and motivational effects. At the psychological level, substances that rapidly reduce anxiety, dampen aversive emotional states, or enhance mood may be particularly attractive for individuals with trauma-related affective dysregulation (Garland et al., 2013). Together, these findings highlight complementary neurobiological and psychological mechanisms that may render opioids and stimulants especially reinforcing in the context of early adversity.

In contrast to nicotine and illicit drug use, alcohol use displayed a more complex pattern. Specifically, ELS exposure was associated with higher likelihood for high-risk drinking and with more severe drinking patterns among drinkers. However, we also found ELS exposure to be associated with increased odds for total abstinence from alcohol, suggesting a polarization towards extreme drinking patterns. Potentially, this polarization reflects differential coping mechanisms in individuals with early adversity. Some individuals may use alcohol for affect regulation or stress relief (Garland et al., 2013), which is supported by findings from our mediation analysis showing that depressive and anxiety symptoms exhibited the largest indirect effect for alcohol-related outcomes. However, others may avoid alcohol entirely, for instance due to fear of losing control. Indeed, anticipated loss of control is one of the central reasons for alcohol abstinence (Epler et al., 2009). In turn, perceived control, i.e. the belief that one can influence outcomes through own actions, has been shown to buffer against the negative impact of ELS on mental health (You et al., 2024), suggesting a potential resilience factor in the context of risky alcohol drinking. Taken together, the simultaneous presence of both abstinence and high-risk drinking highlights heterogeneous behavioral adaptations to early adversity, potentially driven by distinct coping mechanisms.

Using latent class analysis, we identified four distinct substance use profiles, including individuals with minimal or no use, a group primarily characterized by nicotine and cannabis use, a “party drug” class involving predominantly stimulant or club-drug use, and a severe polysubstance group marked by broad multi-drug consumption. ELS exposure was associated with progressively elevated odds of belonging to more severe classes, with the highest risk observed for membership in the severe polysubstance group. This gradient underscores that exposure to ELS not merely increases the likelihood of using individual substances but is associated with more complex and hazardous patterns of multi-drug involvement.

Notably, abuse-related forms of ELS showed the strongest and most consistent associations with substance-related outcomes, whereas neglect-related forms exhibited weaker and more heterogeneous patterns. These findings align with dimensional models of early adversity, positing two partially distinct dimensions of ELS characterized by either threat, involving harm or threat of harm, or deprivation, involving the absence of expected environmental inputs. Threat-related experiences are thought to exert particularly strong effects on neural systems involved in emotional and reward processing (Sheridan & McLaughlin, 2014), thereby increasing vulnerability to risky health behaviors, including substance use (Duffy et al., 2018). Among abuse-related subtypes of ELS, sexual abuse showed the strongest association for most substance use behaviors, including polysubstance use, mirroring prior evidence that survivors of sexual trauma represent a subgroup at particularly elevated risk (Fletcher et al., 2021). These findings underscore the importance of early identification and implementation of tailored, trauma-informed interventions for individuals with histories of sexual abuse, who constitute a priority group for targeted prevention and treatment efforts.

Overall, associations between ELS and later substance use were more pronounced in women than in men, except for nicotine. Evidence suggests that childhood adversity may heighten women’s sensitivity to stress by disrupting adaptive coping and emotion regulation processes, which in turn increase vulnerability to developing substance use disorders in adulthood (Myers et al., 2014; Oldehinkel & Bouma, 2011). Consistent with this idea, studies show that women are more likely than men to use substances as a way to manage distress and regulate negative emotions triggered by traumatic experiences (Brady & Randall, 1999; Hudson et al., 2017). Second, effects were stronger in younger adults. This aligns with evidence showing that neural systems altered by ELS vary in their functional influence across development and are particularly impactful in early adulthood, when stress-sensitive and regulatory neural circuits are still maturing (Teicher et al., 2016). Moreover, younger adults are in developmental periods and psychosocial contexts in which substance use is more normative and opportunities are more prevalent (Chen & Jacobsen, 2012), whereas social and environmental risk decreases with age (Allen et al., 2021), thereby potentially amplifying the impact of ELS in younger individuals.

Several limitations merit consideration. First, ELS exposure was assessed retrospectively via self-report and may be subject to recall bias. Moreover, the Childhood Trauma Screener employed in the present study captures ELS exposure, but not the timing or duration of maltreatment, potentially obscuring more nuanced developmental effects. Future work, particularly smaller-scale studies, should aim to validate ELS by more objective measures, such as official records or reports from multiple informants, and employ instruments that allow more detailed assessments of ELS. Second, although our study assessed substance use and indicators of severity, we lacked information on clinically relevant diagnostic criteria for substance use disorders. As substance use alone does not imply the presence of a disorder, our findings should be interpreted in terms of substance use behavior, rather than clinical diagnoses. Moreover, we could not disentangle medically prescribed use of substances, such as opioids or cannabis, from non-medical use. Third, the cross-sectional study design precludes causal inference. Fourth, residual confounding cannot be ruled out, particularly with regard to early-life socioeconomic conditions as well as familial and genetic factors that may jointly influence exposure to adversity and later substance use. Lastly, future studies should incorporate biological markers relevant to stress and addiction processes, such as markers on HPA-axis functioning (Heim et al., 2008), inflammatory markers (Pace et al., 2006), neuroimaging measures of stress- and reward-related circuitry (Teicher et al., 2016), and epigenetic indicators of stress regulation (Klengel et al., 2013), to further elucidate the mechanisms linking ELS exposure to substance use in adulthood.

In sum, the present analysis demonstrates that ELS exposure is a pervasive risk factor for substance use behaviors in adulthood in Germany. The graded and subtype-specific associations observed across multiple substances and behavioral profiles highlight the enduring influence of childhood adversity on health-relevant behaviors. These findings underscore the need for trauma-informed approaches in clinical practice, public health, and policy to mitigate the long-term consequences of early adversity.

## Contributors

Conceptualization: CH, HT, JB; Methodology: JB, CH; Software: JB; Formal analysis: JB, KMD; Resources: PB, SC, MF, CG, HJG, AK, CL, AL, ML, CMF, DM, RM, KN, NO, AP, TP, MJS, TK, LK; Data curation: PB, SC, MF, CG, HJG, AK, CL, AL, ML, CMF, DM, RM, KN, NO, AP, TP, MJS, TK, LK; Writing—original draft: JB, CH; Writing—review & editing: CH, HT, KMD, AG, AS, PB, SC, MF, CG, HJG, AK, CL, AL, ML, CMF, DM, RM, KN, NO, AP, TP, MJS, TK, LK; Visualization: JB; Supervision: CH, HT, TK, LK. JB and KMD have accessed and verified the data. All authors reviewed the final version of the manuscript and had the responsibility for the decision to submit the paper for publication.

## Data availability statement

Access to data from the German National Cohort Study (“NAKO Gesundheitsstudie”) is subject to approval by the NAKO Use and Access Committee. Qualified researchers may request access to the data through the NAKO application process. For further information is available at https://nako.de/en/research/transferhub/.

## Acknowledgements

This study was supported by the German Research Foundation (Deutsche Forschungsgemeinschaft, DFG TRR 265, Project A04 (C.H. and H.T., grant number 402170461). The current analysis was conducted with data (Application No. NAKO-1111) from the German National Cohort (NAKO) (www.nako.de). The NAKO is funded by the Federal Ministry of Research, Technology and Space (BMFTR, project reference numbers: 01ER1301A/B/C, 01ER1511D, 01ER1801A/B/C/D, and 01ER2301A/B/C), federal states of Germany and the Helmholtz Association, the participating universities and the institutes of the Leibniz Association. We thank all participants who took part in the NAKO study and the staff of this research initiative.

