## Supplementary_Information for "Exposure to childhood adversity and adult alcohol, nicotine, and illicit drug use in Germany: a cross-sectional analysis of the German National Cohort (NAKO)"

**Running Title**: Early Adversity and Substance Use

**TABLE OF CONTENTS**

Supplementary Results: Moderation by age and sex

Table S1. Missing data for all variables, based on the total sample of N = 172,173 subjects.

Table S2. Associations between number of early life stress (ELS) subtypes experienced and substance use, separated by nicotine, alcohol, and illicit drug use.

Table S3. Associations of different early life stress (ELS) subtypes and number of illicit drugs consumed during lifetime.

Table S4. Interaction effects between early life stress (ELS) and age on lifetime illicit drug use.

Table S5. Causal mediation models assessing the indirect effects of depressive and anxiety symptoms on the association between early life stress (ELS) and substance use, separated by nicotine, alcohol, and illicit drug use.

Table S6. Comparison of model fit indices used to determine the optimal latent class solution.

Table S7. Basic characteristics and substance use, stratified for latent substance use classes.

**Supplementary Results: Moderation by age and sex**

*Nicotine*. The association between ELS exposure and lifetime daily cigarette consumption was stronger in men compared to women (β = -0.036, 95% CI [-0.059, -0.012]) and stronger among younger subjects (β = -0.019, 95% CI [-0.032, -0.006]).

*Alcohol*. Sex moderated the associations between ELS exposure and alcohol use outcomes, with stronger effects in women for AUDIT-C total score (β = 0.043, 95% CI [0.019, 0.066]) and drinking frequency (β = 0.035, 95% CI [0.012, 0.059]). Moreover, significant interactions were found with age, showing that ELS exposure was more strongly linked to higher daily alcohol intake among younger individuals (β = –0.024, 95% CI [–0.036, –0.011]), whereas its association with AUDIT-C total score was stronger in older individuals (β = 0.020, 95% CI [0.007, 0.032]).

*Illicit drugs*. Both sex and age significantly moderated the association between ELS exposure and substance use. Specifically, the association between ELS and the use of cannabis (OR = 1.22, 95% CI [1.16, 1.30]), amphetamine-type stimulants (OR = 1.30, 95% CI [1.15, 1.46]), LSD (OR = 1.25, 95% CI [1.09, 1.43]), non-heroin opioids (OR = 1.25, 95% CI [1.06, 1.47]), and powder cocaine (OR = 1.29, 95% CI [1.16, 1.45]) was stronger in women compared to men. Moreover, stronger associations between ELS exposure and illicit drug use were observed among younger individuals for all substances, except for synthetic cannabinoids (Table S4).

Table S1. Missing data for all variables, based on the total sample of N = 172,173 subjects.

| **Variable** | **% missing of total sample** |
| --- | --- |
| **Demographics** |  |
| Sex | 0.00 |
| Age | 0.00 |
| Site | 0.00 |
| ISCED-97 years of education | 2.43 |
| Parental employment | 5.21 |
| Parental education | 16.38 |
| **Early life stress (ELS)** |  |
| CTS: ELS exposure (yes/no) | 0.00 |
| CTS: Total score | 0.08 |
| CTS: Physical neglect | 0.06 |
| CTS: Emotional neglect | 0.01 |
| CTS: Physical abuse | 0.03 |
| CTS: Emotional abuse | 0.04 |
| CTS: Sexual abuse | 0.04 |
| **Clinical characteristics** |  |
| Depression (PHQ-9 total score) | 0.62 |
| Anxiety (GAD-7 total score) | 0.65 |
| **Alcohol use** |  |
| AUDIT-C total score | 0.16 |
| AUDIT-C item 1 | 0.14 |
| AUDIT-C item 2 | 7.93 |
| AUDIT-C item 3 | 7.93 |
| Average alcohol intake (g/day) | 0.48 |
| **Nicotine use** |  |
| Lifetime smoking prevalence | 0.18 |
| Lifetime smoking duration | 0.28 |
| Lifetime daily cigarette consumption | 0.19 |
| Age at smoking initiation | 47.25 |
| **Illicit drug use** |  |
| Cannabis | 2.40 |
| Synthetic cannabinoids | 2.55 |
| Amphetamine-type stimulants | 2.50 |
| Methamphetamine | 2.54 |

| MDMA | 2.54 |
| --- | --- |
| LSD | 2.56 |
| Heroin | 2.55 |
| Non-heroin opioids | 2.56 |
| Powder cocaine | 2.56 |
| Crack cocaine | 2.55 |

Table S1. Missing data for all variables, based on the total sample of N = 172,173 subjects (continued).

Abbreviations: AUDIT-C = Alcohol Use Disorder Identification Test – Consumption; CTS = Childhood Trauma Screener; ELS = Early life stress; GAD-7 = Generalized Anxiety Disorder Scale; ISCED-97 = International Standard Classification of Education 97; PHQ-9 = Patient Health Questionnaire-9

Table S2. Associations between number of early life stress (ELS) subtypes experienced and substance use, separated by nicotine, alcohol, and illicit drug use.

| **Substance** | **Outcome** | **Effect estimate** | **95% CI** |
| --- | --- | --- | --- |
| Nicotine | Lifetime smoking prevalence | OR = 1.26 | [1.25, 1.28] |
|  | Lifetime smoking duration | β = 0.081 | [0v076, 0.086] |
|  | Lifetime daily cigarette consumption | β = 0.090 | [0.085, 0.095] |
|  | Age at smoking initiation | β = −0.018 | [−0.025, −0.011] |
| Alcohol | Hazardous drinking (AUDIT-C total score) | β = −0.028 | [−0.033, −0.023] |
|  | Drinking frequency (AUDIT-C item 1) | β = −0.049 | [−0.054, −0.044] |
|  | Number of drinks per occasion (AUDIT-C item 2) | β = 0.020 | [0.014, 0.025] |
|  | Binge-drinking frequency (AUDIT-C item 3) | β = 0.008 | [0.003, 0.013] |
|  | Average amount of daily alcohol intake | β = 0.003 | [−0.002, 0.008] |
| Illicit drugs | Cannabis | OR = 1.25 | [1.23, 1.27] |
|  | Synthetic cannabinoids | OR = 1.37 | [1.25, 1.50] |
|  | Amphetamine-type stimulants | OR = 1.42 | [1.38, 1.47] |
|  | Methamphetamine | OR = 1.56 | [1.43, 1.70] |
|  | MDMA | OR = 1.43 | [1.38, 1.48] |
|  | LSD | OR = 1.44 | [1.39, 1.49] |
|  | Heroin | OR = 1.56 | [1.45, 1.67] |
|  | Non-heroin opioids | OR = 1.49 | [1.43, 1.55] |
|  | Powder cocaine | OR = 1.40 | [1.36, 1.44] |
|  | Crack cocaine | OR = 1.47 | [1.31, 1.64] |

Abbreviations: AUDIT-C = Alcohol Use Disorder Identification Test – Consumption; β = standardized beta coefficient; CI = confidence interval; OR = Odds Ratio

Table S3. Associations of different early life stress (ELS) subtypes and number of illicit drugs consumed during lifetime.

| **ELS subtype** | **Standardized beta coefficient** | **95% CI** |
| --- | --- | --- |
| Emotional neglect | 0.089 | [0.066, 0.111] |
| Physical neglect | 0.118 | [0.096, 0.140] |
| Emotional abuse | 0.202 | [0.180, 0.224] |
| Sexual abuse | 0.244 | [0.221, 0.267] |
| Physical abuse | −0.067 | [−0.085, −0.049] |

Abbreviations: CI = confidence interval; ELS = Early life stress

Table S4. Interaction effects between early life stress (ELS) and age on lifetime illicit drug use.

| **Substance** | **OR** | **95% CI** |
| --- | --- | --- |
| Cannabis | 0.97 | [0.95, 0.99] |
| Synthetic cannabinoids | 1.01 | [0.88, 1.16] |
| Amphetamine-type stimulants | 0.91 | [0.87, 0.96] |
| Methamphetamine | 0.82 | [0.70, 0.95] |
| MDMA | 0.85 | [0.81, 0.90] |
| LSD | 0.94 | [0.89, 0.99] |
| Heroin | 0.81 | [0.71, 0.93] |
| Non-heroin opioids | 0.85 | [0.79, 0.90] |
| Powder cocaine | 0.87 | [0.84, 0.91] |
| Crack cocaine | 0.72 | [0.58, 0.89] |

Abbreviations: CI = confidence interval; OR = Odds Ratio

Table S5. Causal mediation models assessing the indirect effects of depressive and anxiety symptoms on the association between early life stress (ELS) and substance use, separated by nicotine, alcohol, and illicit drug use.

| **Substance** | **Outcome** | **Depressive symptoms** | | **Anxiety symptoms** | |
| --- | --- | --- | --- | --- | --- |
|  |  | **Indirect mediation effect in %** | **95% CI** | **Indirect mediation effect in %** | **95% CI** |
| Nicotine | Lifetime smoking prevalence | 12.02 | [11.00, 12.58] | 9.38 | [8.81, 11.25] |
|  | Lifetime smoking duration | 14.00 | [13.73, 15.85] | 9.57 | [8.37, 10.45] |
|  | Lifetime daily cigarette consumption | 13.20 | [12.17, 13.96] | 9.26 | [7.85, 10.25] |
|  | Age at smoking initiation | 10.85 | [2.81, 23.30] | 5.18 | [1.33, 57.87] |
| Alcohol | Drinking frequency  (AUDIT-C item 1) | 13.93 | [12.75, 16.39] | 0.34 | [-0.95, 1.15] |
|  | Number of drinks per occasion (AUDIT-C item 2) | 58.25 | [51.78, 83.76] | 29.00 | [20.76, 50.48] |
|  | Binge-drinking frequency (AUDIT-C item 3) | 62.70 | [35.71, 86.30] | 49.28 | [31.24, 72.54] |
| Illicit drugs | Cannabis | 16.15 | [14.76, 17.30] | 12.49 | [11.40, 13.72] |
|  | Synthetic cannabinoids | 19.18 | [14.99, 31.03] | 14.89 | [11.77, 22.80] |
|  | Amphetamine-type stimulants | 14.20 | [13.36, 17.31] | 10.71 | [10.00, 12.12] |
|  | Methamphetamine | 12.64 | [10.73, 22.75] | 9.93 | [5.76, 13.73] |
|  | MDMA | 12.51 | [12.07, 14.25] | 10.87 | [10.02, 13.01] |
|  | LSD | 11.44 | [10.39, 14.88] | 8.39 | [7.55, 9.94] |
|  | Heroin | 14.45 | [11.81, 18.67] | 11.47 | [9.66, 15.35] |
|  | Non-heroin opioids | 20.46 | [18.10, 22.81] | 15.34 | [13.37, 15.49] |
|  | Powder cocaine | 12.96 | [9.84, 14.42] | 10.97 | [10.23, 12.38] |
|  | Crack cocaine | 16.19 | [6.69, 36.07] | 11.84 | [6.11, 14.52] |

Abbreviations: AUDIT-C = Alcohol Use Disorder Identification Test – Consumption; CI = confidence interval

Table S6. Comparison of model fit indices used to determine the optimal latent class solution.

| **Number of latent classes** | **Bayesian Information Criterion (BIC)** | **Difference in BIC to previous model** | **Akaike Information Criterion (AIC)** | **Log-Likelihood** | **Entropy** |
| --- | --- | --- | --- | --- | --- |
| 1 | 798,591.0 | - | 798,470.8 | -399,223.4 | Inf |
| 2 | 685,276.0 | 113,315.0 | 685,025.4 | -342,487.7 | 0.94 |
| 3 | 670,787.3 | 14,488.7 | 670,406.5 | -335,165.2 | 0.81 |
| 4 | 668,509.4 | 2,277.9 | 667,998.3 | -333,948.2 | 0.67 |
| 5 | 668,721.9 | -212.5 | 668,080.5 | -333,976.3 | 0.66 |
| 6 | 666,666.4 | 2,055.5 | 665,894.8 | -332,870.4 | 0.61 |
| 7 | 665,863.9 | 802.5 | 664,961.9 | -332,390.9 | 0.64 |
| 8 | 665,745.6 | 118.3 | 664,713.3 | -332,253.7 | 0.66 |

Table S7. Basic characteristics and substance use, stratified for latent substance use classes.

| **Variable** | | **Class 1: Minimal substance use**  **(N = 125,110)** | **Class 2: Cannabis**–**nicotine use  (N = 33,772)** | **Class 3: Stimulant**–**hallucinogen use (N = 6,804)** | **Class 4:  High-risk** **poly-substance use (N = 648)** |
| --- | --- | --- | --- | --- | --- |
| **Early life stress (ELS)** | |  |  |  |  |
|  | CTS: ELS exposure  (% yes) | 25.11 | 28.03 | 31.41 | 45.37 |
|  | CTS: Total score | 7.19 (± 2.62) | 7.38 (± 2.79) | 7.63 (± 3.02) | 8.78 (± 3.48) |
|  | CTS: Physical neglect (% yes) | 10.55 | 7.80 | 6.64 | 12.19 |
|  | CTS: Emotional neglect (% yes) | 7.21 | 8.03 | 9.59 | 18.36 |
|  | CTS: Physical abuse  (% yes) | 7.42 | 9.83 | 12.01 | 18.83 |
|  | CTS: Emotional abuse (% yes) | 7.09 | 10.50 | 14.67 | 21.45 |
|  | CTS: Sexual abuse  (% yes) | 5.44 | 7.88 | 9.49 | 11.27 |
| **Demographics** | |  |  |  |  |
|  | Sex (% female) | 52.36 | 43.44 | 33.01 | 25.15 |
|  | Age (years) | 50.30 (± 12.56) | 45.11 (± 12.02) | 40.38 (± 11.51) | 46.24 (± 12.33) |
|  | ISCED-97 years of education | 15.46 (± 2.34) | 15.76 (± 2.29) | 15.39 (± 2.36) | 14.22 (± 2.64) |
|  | Parental employment (range: 0-2) | 1.59 (± 0.53) | 1.61 (± 0.52) | 1.66 (± 0.51) | 1.59 (± 0.53) |
|  | Parental education (range: 2-10) | 5.53 (± 2.03) | 6.05 (± 2.12) | 6.48 (± 2.14) | 5.83 (± 2.18) |
| **Self-reported mental health** | | |  |  |  |
|  | Depressive symptoms (PHQ-9 total score) | 3.68 (± 3v59) | 4.29 (± 3.85) | 5.02 (± 4.34) | 6.17 (± 5.44) |
|  | Anxiety symptoms (GAD-7 total score) | 3.01 (± 3.14) | 3.48 (± 3.32) | 4.00 (± 3.62) | 4.51 (± 4.35) |
| **Nicotine use** | |  |  |  |  |
|  | Lifetime smoking occurrence | 39.06 | 94.50 | 87.64 | 89.66 |
|  | Lifetime smoking duration | 8.00 (± 13.03) | 17.37 (± 11.94) | 16.26 (± 11.67) | 21.44 (± 13.64) |
|  | Lifetime daily cigarette consumption | 5.10 (± 9.04) | 11.60 (± 9.73) | 10.94 (± 9.66) | 13.51 (± 10.56) |
|  | Age at smoking initiation | 18.58 (± 6.72) | 17.46 (± 5.19) | 16.65 (± 4.59) | 15.98 (± 5.49) |
| **Alcohol use** | |  |  |  |  |
|  | Hazardous drinking  (AUDIT-C total score) | 3.15 (± 1.96) | 4.15 (± 2.16) | 4.57 (± 2.33) | 4.11 (± 2.85) |
|  | Drinking frequency  (AUDIT-C item 1) | 2.21 (± 1.18) | 2.55 (± 1.13) | 2.53 (± 1.14) | 2.27 (± 1.37) |
|  | Number of drinks per occasion (AUDIT-C item 2) | 1.28 (± 0.57) | 1.54 (± 0.81) | 1.76 (± 0.96) | 1.83 (± 1.01) |
|  | Binge-drinking frequency  (AUDIT-C item 3) | 0.75 (± 0.85) | 1.14 (± 0.93) | 1.40 (± 0.97) | 1.36 (± 1.11) |
|  | Average amount of daily alcohol intake (g/day) | 9.01 (± 16.41) | 13.63 (± 19.83) | 16.14 (± 23.12) | 17.99 (± 28.81) |
| Table S7. Basic characteristics and substance use, stratified for latent substance use classes (continued). | | | | | |
| **Illicit drug use** | |  |  |  |  |
|  | Cannabis (% yes) | 9.55 | 98.92 | 98.90 | 99.85 |
|  | Synthetic cannabinoids (% yes) | < 0.01 | 0.87 | 5.95 | 44.75 |
|  | Amphetamine-type stimulants (% yes) | 0.28 | 4.74 | 73.94 | 96.30 |
|  | Methamphetamine  (% yes) | 0.00 | 0.06 | 5.44 | 42.28 |
|  | MDMA (% yes) | 0.04 | 2.50 | 70.71 | 85.65 |
|  | LSD (% yes) | 0.00 | 3.84 | 50.90 | 94.60 |
|  | Heroin (% yes) | < 0.01 | 0.12 | 4.38 | 78.40 |
|  | Non-heroin opioids  (% yes) | 0.66 | 2.84 | 13.93 | 94.29 |
|  | Powder cocaine (% yes) | 0.07 | 7.01 | 80.78 | 99.07 |
|  | Crack cocaine (% yes) | 0.01 | 0.10 | 0.90 | 36.88 |

Abbreviations: AUDIT-C = Alcohol Use Disorder Identification Test – Consumption; CTS = Childhood Trauma Screener; ELS = Early life stress; GAD-7 = Generalized Anxiety Disorder Scale; ISCED-97 = International Standard Classification of Education 97; PHQ-9 = Patient Health Questionnaire-9
